# Quantitative longitudinal predictions of Alzheimer’s disease by multi-modal predictive learning

**DOI:** 10.1101/2020.06.04.133645

**Authors:** M. Prakash, M. Abdelaziz, L. Zhang, B.A. Strange, J. Tohka, Alzheimer’s Disease Neuroimaging Initiative

**Affiliations:** University of Eastern Finland, A.I. Virtanen Institute for Molecular Sciences, Kuopio, Finland; Zewail City of Science and Technology, Giza, Egypt; Department of Neuroimaging, Alzheimer’s Disease Research Centre, Reina Sofia-CIEN Foundation, Madrid, Spain; Laboratory for Clinical Neuroscience, CTB, Universidad Politécnica de Madrid, Madrid, Spain

**Author notes:** **Corresponding Author**: Mithilesh Prakash, PhD, A.I. Virtanen Institute for Molecular Sciences, University of Eastern Finland, P.O.B. 1627, FI-70211 Kuopio, Finland. Data used in the preparation of this article were obtained from the Alzheimer’s Disease Neuroimaging Initiative (ADNI) database.

**Keywords:** Alzheimer’s disease, Magnetic resonance imaging, Machine Learning, Neuropsychology, Multivariate, PLS, ADAS-cog

## Abstract

**Background:** Quantitatively predicting the progression of Alzheimer’s disease (AD) in an individual on a continuous scale, such as AD assessment scale-cognitive (ADAS-cog) scores, is informative for a personalized approach as opposed to qualitatively classifying the individual into a broad disease category. We hypothesize that multi-modal data and predictive learning models can be employed for longitudinally predicting ADAS-cog scores.

**Methods:** Multivariate regression techniques were employed to model baseline multi-modal data (demographics, neuroimaging, and cerebrospinal fluid based markers, and genetic factors) and future ADAS-cog scores. Prediction models were subjected to repeated cross-validation and the resulting mean absolute error and cross-validated correlation of the model assessed.

**Results:** Prediction models on multi-modal data outperformed single modal data up to 36 months. Incorporating baseline ADAS-cog scores to prediction models marginally improved predictive performance.

**Conclusions:** Future ADAS-cog scores were successfully estimated via predictive learning aiding clinicians in identifying those at greater risk of decline and apply interventions at an earlier disease stage and inform likely future disease progression in individuals enrolled in AD clinical trials.

## 1 Background

Alzheimer’s disease (AD) is an irreversible and multi-factorial neurodegenerative disease with a progressive decline in cognitive abilities [1]. AD affects several tens of millions of people globally. Yet, the pathogenesis of AD remains unclear [2]. Cognitive tests, brain volumetry from magnetic resonance imaging (MRI), amyloid load and glucose consumption levels from positron emission tomography (PET), and protein analysis of cerebrospinal fluid (CSF) provide valuable and complementary disease markers to chart the disease progression [3]. Qualitative manual analysis of these markers to diagnose patients could be potentially aided by automated algorithms.

The classification based on clinical diagnosis places an individual into normal, mild cognitive impairment (MCI), or AD groups [4]. Memory loss (either self-reported or by an associate) is observed during the initial stages of AD [5]. Declining cognitive skills is also common and can potentially lead to dementia [6]. Hence, it is imperative that the disease progression is carefully monitored at the earliest stages [7]. AD risk factors include sociodemographic factors (e.g., increasing age and fewer years of education), genetic (APOE expression) and patient medical and family history [8]. A clinical diagnosis of AD is currently a challenge due to lack of clear diagnostic markers of AD, and overlapping clinical features with other dementia types. However at postmortem, AD is characterized by the presence of amyloid β-peptide plaques and accumulations of **τ** proteins in the brain histology samples [9].

The progressive nature of AD makes diagnosing an individual into any of the discrete groups a challenging proposition [10,11]. Conventional progression tracking analyzes clinical changes in MRI, CSF and cognitive biomarkers [12,13], but this could be inefficient as the changes can be slow and difficult to detect [14,15]. The change in these biomarkers is nonlinear with AD’s progression, further complicating longitudinal tracking. Therefore, quantifying and tracking the condition of the patient by continuous measures such as ADAS-cog scores has been advocated [16,17]. ADAS-cog is widely used clinically (to measure language, memory, praxis, and other cognitive abilities) and provides an accurate description of the cognitive state on a continuous scale, making it an ideal choice in our study [18,19]. The availability of standardized multi-modal data and corresponding longitudinal ADAS-cog scores from research organizations, such as the Alzheimer’s Disease Neuroimaging Initiative (ADNI) project, has enabled the development of novel techniques for tracking AD progression by employing machine learning [20]. However, predicting ADAS-cog scores has been reported as very difficult [21]. In the recent Alzheimer’s Disease Prediction of Longitudinal Evolution (TADPOLE) Challenge (https://tadpole.grand-challenge.org/), forecasts of clinical diagnosis and ventricle volume were very good, whereas, for ADAS-cog, no team participating in the challenge was able to generate forecasts that were significantly better than chance.

Multivariate regression techniques, such as partial least squares regression (PLSR), support-vector regression (SVR) and random forest regression, enable modeling complex relationships between baseline multi-modal ADNI data (predictors) with future ADAS-cog 13 scores [22,23]. The multivariate nature of the modeling is desirable for the ADAS-cog score trajectory analysis due to the complementary nature of the AD measures. The resulting trajectory predictions could alert clinicians to prescribe appropriately (once disease modifying interventions are available). Moreover, knowing the likely future trajectory of the disease will provide a benchmark with which to test clinical evolution in patients enrolled in clinical trials.

We hypothesized that the multivariate regression techniques are well suited for multi-factorial diseases and that the progression of AD, as indicated by ADAS-cog scores in subsequent timelines, can be accurately predicted. Furthermore, the inclusion of baseline ADAS-cog scores could improve the predictions of the model in subsequent follow-ups.

## 2 Methods

### 2.1 ADNI Dataset

Data in this study were obtained from the Alzheimer’s Disease Neuroimaging Initiative (ADNI) database (http://adni.loni.usc.edu/). In addition to the various summary tables directly provided by ADNI, we used summary tables prepared for the TADPOLE grand challenge based on ADNI data at https://tadpole.grand-challenge) [21,24]. The data are from the TADPOLE tables if not otherwise stated. Specific variable names are provided as supplementary Table S.1. The ADNI project started in 2003 as a public-private partnership, led by PI Michael W. Weiner, MD. The main objective of ADNI is to evaluate the application of serial magnetic resonance imaging (MRI), positron emission tomography (PET), other biological markers, and clinical and neuropsychological assessment in a multi-modal approach to determine the longitudinal progression of mild cognitive impairment (MCI) and early Alzheimer’s disease (AD). We utilized pre-processed ADNI data because of the standardized processing pipeline that ensured the quality of the data. This multimodal data is readily available for other researchers enabling a direct comparison of the study results. Readers are directed to www.adni-info.org for detailed information on the ADNI project and the TADPOLE challenge https://tadpole.grand-challenge.org constructed by the EuroPOND consortium (http://europond.eu).

#### 2.1.1 Subjects

The characteristics of subjects recruited in the ADNI dataset are described in detail here http://adni.loni.usc.edu/. The trends of the ADAS-cog 13 scores utilized in this study are provided in Figure S.1 and the details of subject characteristics are provided in Table S.2 of the supplementary section. There are fewer subjects in follow-up visits than in the baseline visit due to subject attrition and missing data. Note that some subjects change diagnostic status over the followup period. The roster identification (RID) numbers of the included subjects are provided as comma-separated values in the supplementary section.

#### 2.1.2 MRI

As MRI features, we used 9 features: intracranial volume (ICV), and volumes of the hippocampus, entorhinal cortex, and lateral ventricles as well as the latter four divided by the ICV. These features were selected based on previous studies [25]. We included volumes divided by the ICV as it is unclear whether raw or ICV-corrected volumes are better predictors of dementia [25,26]. MR imaging protocol details are provided by ADNI at http://adni.loni.usc.edu/methods/mri-tool/mri-analysis/. Cortical reconstruction and volumetric segmentation had been performed with the FreeSurfer 5.1 image analysis suite. A brief description of the processing is provided in the supplementary material (Section B) [27].

#### 2.1.3 AV-45 PET

As AV-45 PET features, we used standardized uptake values (SUVs) in four regions: frontal cortex, cingulate, lateral parietal cortex, and lateral temporal cortex. The AV-45 PET measures amyloidbeta load in the brain. AV-45 PET imaging and preprocessing details are available at http://adni.loni.usc.edu/methods/pet-analysis-method/pet-analysis/ [28]. We used regional SUV ratios processed according to the UC Berkeley protocol [28-30]. Each AV-45 PET scan was coregistered to the corresponding MRI and the mean AV-45 uptake within the regions of interest and reference regions was calculated. Regions of interest were composites of frontal regions, anterior/posterior cingulate regions, lateral parietal regions, and lateral temporal regions [31]. The final PET measurements were the average amyloid-beta uptakes in the four ROIs normalized by the whole cerebellum reference region.

#### 2.1.4 FDG PET

As FDG-PET features, we used average SUVs in five brain regions: bilateral angular gyri, bilateral posterior cingulate gyri, and bilateral inferior temporal gyri. The FDG PET data measures glucose consumption and is shown to be strongly related to dementia and cognitive impairment when compared to normal control subjects [30,32,33]. Motion correction and co-registration with MRI was performed on the acquired PET data. The highest 50% of voxel values within a hand-drawn pons/cerebellar vermis region were selected and their mean was used to normalize each ROI measurement resulting in the final FDG PET measurements. Regions of interests were bilateral angular gyri, bilateral posterior cingulate gyri, and bilateral inferior temporal gyri.

#### 2.1.5 CSF proteins

The baseline CSF Aβ42, t-tau, and p-tau were used as CSF features [34]. CSF was collected in the morning after an overnight fast using a 20- or 24-gauge spinal needle, frozen within 1 hour of collection, and transported on dry ice to the ADNI Biomarker Core laboratory at the University of Pennsylvania Medical Center. The levels of Aβ42, t-tau, and p-tau in CSF were used.

#### 2.1.6 Neuropsychology and behavioral (NePB) assessments

The NePB assessments reflect the cognitive abilities of the subjects. Subjects underwent a battery of NePB tests [35]. We selected to include 5 NePB scores as NePB features: the summary score from Mini-Mental State Examination (MMSE) [36], three summary scores of Rey’s auditory verbal learning test (RAVLT; learning, immediate, and percent forgetting) [37], and a summary score from the functional activities Questionnaire (FAQ) [38].

#### 2.1.7 Risk factors: age, education, and APOE

Past studies have found several risk factors contributing to AD [8]. We considered age, the number of APOE e4 alleles, and the years of education. With aging, normal cognitive decline is an accepted phenomenon, but lower education and lower cerebral metabolic activity could accelerate the normal decline [39]. The APOE e4 allele, present in approximately 10-15% of people, increases the risk for late-onset AD and lowers the age of onset. One copy of e4 (e3/e4) can increase risk by 2-3 times while homozygotes (e4/e4) can be at 12 times increased risk [40]. We coded APOE e4 status of absence, single copy or homozygous coded as 0, 1 and 2 respectively.

#### 2.1.8 ADAS-cog scores

The ADAS-cog 11 task scale was developed to assess the efficacy of anti-dementia treatments. Further developments to the scale shifted its sensitivity towards pre-dementia syndromes as well, primarily mild cognitive impairment (MCI). The ADAS-cog 13 task scale was one such improvement on the original ADAS-cog 11, with additional memory and attention/executive function tasks [41]. The final 13 tasks test verbal memory (3 tasks), clinician-rated perception (4 tasks), and general cognition (6 tasks). It was found to perform better than the ADAS-cog 11 at discriminating between MCI and mild AD patients, as well as have better sensitivity to treatment effects in MCI [42]. As the ADAS-cog 13 fully encompasses the ADAS-cog 11 tasks, it is also backward compatible. As such, we used the ADAS-cog 13 scale for our study as a continuous quantitative measure of a subject’s disease status. The scores at baseline, 12-month, 24-month and 36-month timelines were obtained from the ADNI dataset (Table S.2). The value (0 to 85) of these scores is lowest for the normal control group and increases with disease progression and the scores are highest for AD subjects.

### 2.2 Multivariate regression analysis

We employed multivariate regression to predict ADAS-cog scores based on predictor variables detailed in section 2.1. We considered four different prediction tasks: predicting ADAS-cog score at baseline and at 12, 24, or 36 months after the baseline. In all of these tasks, the predictor variables are from the baseline visit. The group of features (predictors) used for regression are denoted by the column vectors X_i_, (i = 0, 1,…, L), where L is the number of features (Figure S.2). The ADAS-cog scores (dependent variable or response variable) are denoted by the column vector Y.

We employed widely used machine learning techniques including partial least squares regression (PLSR) [43], support vector regression (SVR) [44], and random forest regression (RF) and created prediction models [45]. Additionally, a genetic algorithm (GA) was utilized to rank the variables in the order of importance in the multi-modal case [46]. The details on these methods are provided in the supplementary section.

### 2.3 Regression modeling and performance metrics

The prediction of the ADAS scores (at baseline, 12-months, 24-months, and 36 months) was performed by employing PLSR, SVR, and RF. Both single modal (each modality of Section 2.1 alone) and multi-modal predictors (all modalities of Section 2.1 combined,) were considered. All the predictors were from the baseline visit. We evaluated the prediction models using 5-fold repeated cross-validation with 10 repeats, see **Figure 1** and Figure S.2. Under single modalities Age, years of education (Edu), number of APOE e4 alleles (APOE) were exactly 1 variable each, CSF had 3 variables, AVF45-PET had 4 variables, NePB and FDG had 5 variables each, MRI had 9 variables and hence the multimodal model had a total of 29 variables (Figure S.2). All variables were assumed to be continuous and we standardized the variables to be zero-mean and unit standard deviation. The model was evaluated in terms of correlation coefficient (***ρ***) and the mean absolute error (MAE) between the actual ADAS-cog 13 scores and its model-predicted values. From the 5fold cross-validation, we averaged the resulting 5 distinct values and computed 95% confidence intervals (CIs) using the bootstrapping method. Similarly, MAE and its CIs were computed. The process was repeated 10 times and its distribution analyzed. For mathematical details of these performance metrics as well as the CI computation in the case of repeated cross-validation that takes into account inter-dependency of distinct repeats, readers may consult Lewis et.al. [47].

**Figure 1:**
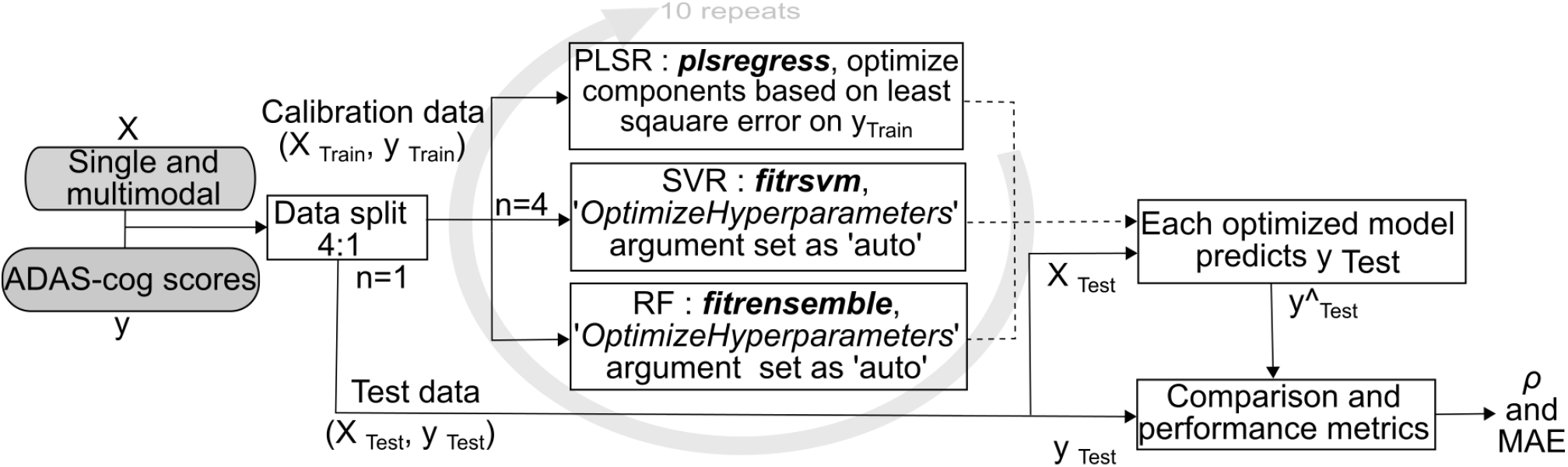
Schematic of regression modeling. X is single or multi-modal predictors and Y is the target value to be predicted. We utilized 5-fold cross-validation repeated 10 times to account for the random assignment of subjects to different folds. Partial least squares regression (PLSR), supportvector regression (SVR) and random forest regression (RF) models were trained and tuned based on training folds and evaluated on test folds. The utilized Matlab function and hyperparameter tuning are shown in italics. Cross-validated correlation (***ρ***) and mean absolute error (MAE) metrics were employed and average performance for 10 runs computed.

The analyses were performed on MATLAB 2018b (The Mathworks Inc, Natick, MA) using native machine learning functions. The PLSR was executed with *plsregress* function and the optimal number of PLS components was manually selected based on the least root mean square error for training data [48]. SVR was executed with *fitrsvm* and RF with *fitrensemble* and in both methods the models were tuned by setting *OptimizeHyperparameters* argument as *auto* [49,50]. Additionally, Additionally, GA-PLS was utilized to analyze the importance of each modality in the multimodal PLSR regression models [51].

(The main codes and resulting *.mat* file are available on GitHub: https://github.com/mithp/ADAS_multimodal.git)

## 3 Results

As depicted in **Figure 1**, we created single modality and multi-modal regression models and compared their performance. The comparison (**Figure 3**) shows that multi-modal based prediction models outperform single modality consistently in all the timelines (baseline and subsequent 12, 24 and 36-month follow-up) in all subjects tested (i.e., collapsing over diagnostic categories). The correlation between the predicted ADAS-Cog 13 based on multi-modal data and that observed at 12, 24 and 36 months, reached 0.86, 0.82, and 0.75, respectively. The performance comparison (Figure S.3) shows that the differences among PLSR, SVR, and RF were not significant (i.e., p > 0.05), except for some instances where PLSR underperformed compared to RF (baseline and 12 months: MRI, CSF, and FDG; 24 and 36 months: APOE and multi-modal). However, PLSR models were computationally faster and performed consistently.

By analyzing the importance of measures (**Figure 2**) contributing to PLSR’s correlation we observe that the neuropsychological and behavioral parameters (NePB) were most important and consistent across time periods for predicting ADAS score, followed by CSF and MRI biomarkers. Despite the association of age at baseline, years of education (Edu.) and APOE e4 status with AD risk, thers parameters were found to be least important, perhaps because these factors are somehow reflected in other parameters. By contrast, the importance of amyloid and τ increased when predictions were made 36 months in advance (**Figure 2**). Additionally, metabolic activity in temporal right and left sides were on the opposite ends of the importance in the ADAS-cog score predictions.

**Figure 2:**
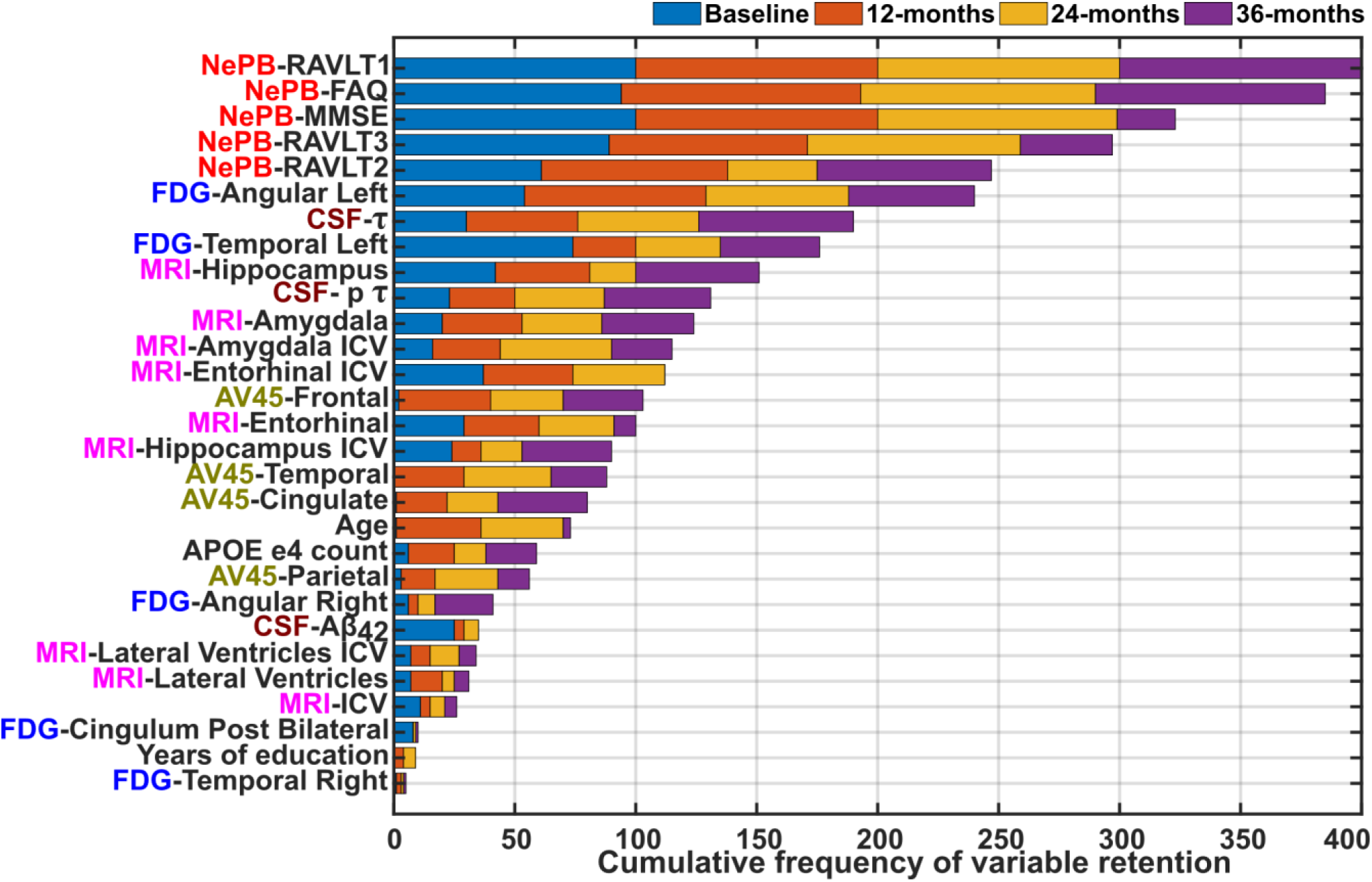
Genetic algorithm-based importance of parameters in correlations as observed for 100 runs for every time period. The frequency indicates the proportional contribution in ADAS-cog 13 score prediction. The modality group is prefixed to the variable names.

Grouping data based on diagnosis at baseline (**Figure 4**) and analyzing the performance further magnified the poor correlation when a single modality approach was employed to predict this multi-factorial disease. We observe that NePB, single modal, data shows the best predictive performance, in keeping with the fact that the to-be-predicted variable (ADAS-Cog 13) also contains NeBP outcomes. However, the multimodal approach performs better than MCI and AD groups especially during 24- and 36-month time periods. Due to the high variation in ADAS scores in AD groups the correlation (***ρ***) and MAE were not inversely proportional to each other.

**Figure 3:**
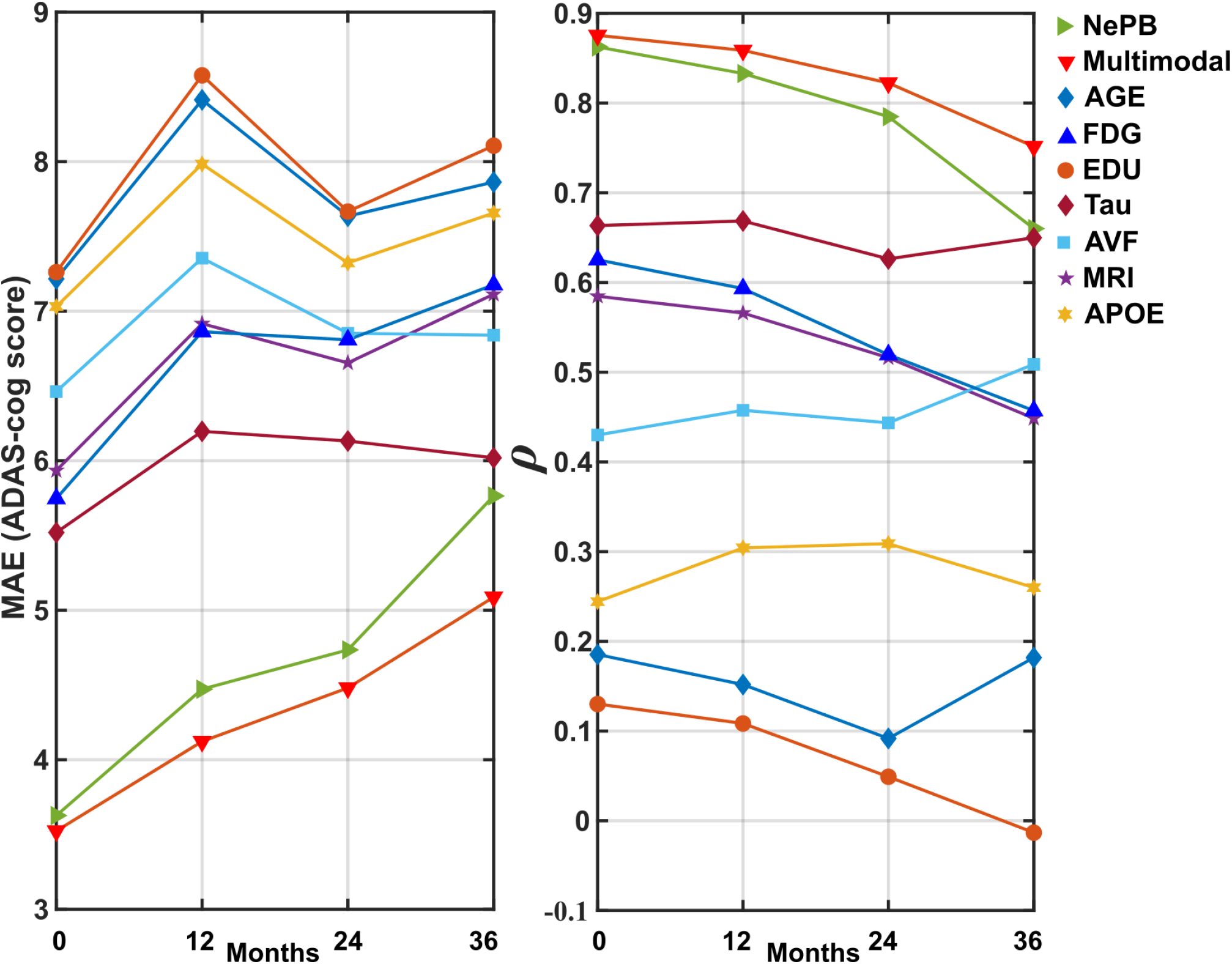
Comparison of single and multi-modal dataset performance - collapsing across diagnostic status - with partial least squares regression. The performance measures cross-validation correlation (***ρ***) and mean absolute error (MAE) for ADAS-cog scores are plotted for predictions at 0, 12, 24 and 36 months in advance.

**Figure 4:**
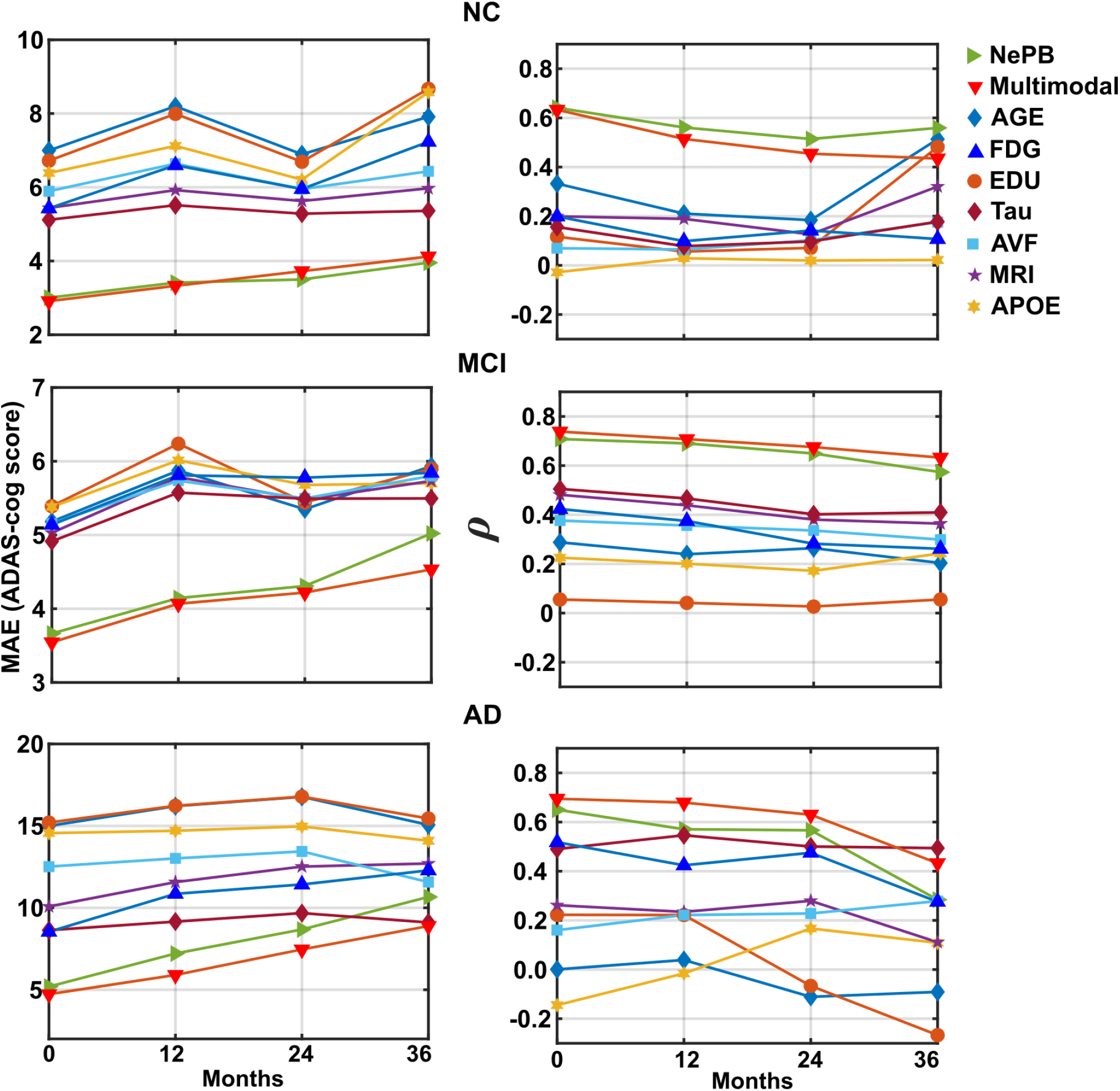
Performance of PLSR on single- and multi-modal data stratified according to baseline clinical diagnosis [normal cognition (**NC**), mild cognitive impairment (**MCI**) and Alzheimer’s disease (**AD**)]. The performance measures cross-validation correlation (***p***) and mean absolute error (MAE) for ADAS-cog scores are plotted for predictions at 0, 12, 24 and 36 months in advance.

Our multimodal approach (multivariate) based prediction models with the inclusion of baseline ADAS-cog scores were better (***ρ*** = 0.80 to 0.90, **Figure 5**) than prediction models based only on baseline ADAS-cog scores (univariate, ***ρ*** = 0.75 to 0.87). The inclusion of the ADAS-cog score with other baseline multi-modal predictors was observed with improvements (p = 0.002 to 0.18) in the correlations. Overall, the prediction models predict well across the time periods and this can be observed when we compare the mean predicted values versus the actual mean values (**Figure 6**).

**Figure 5:**
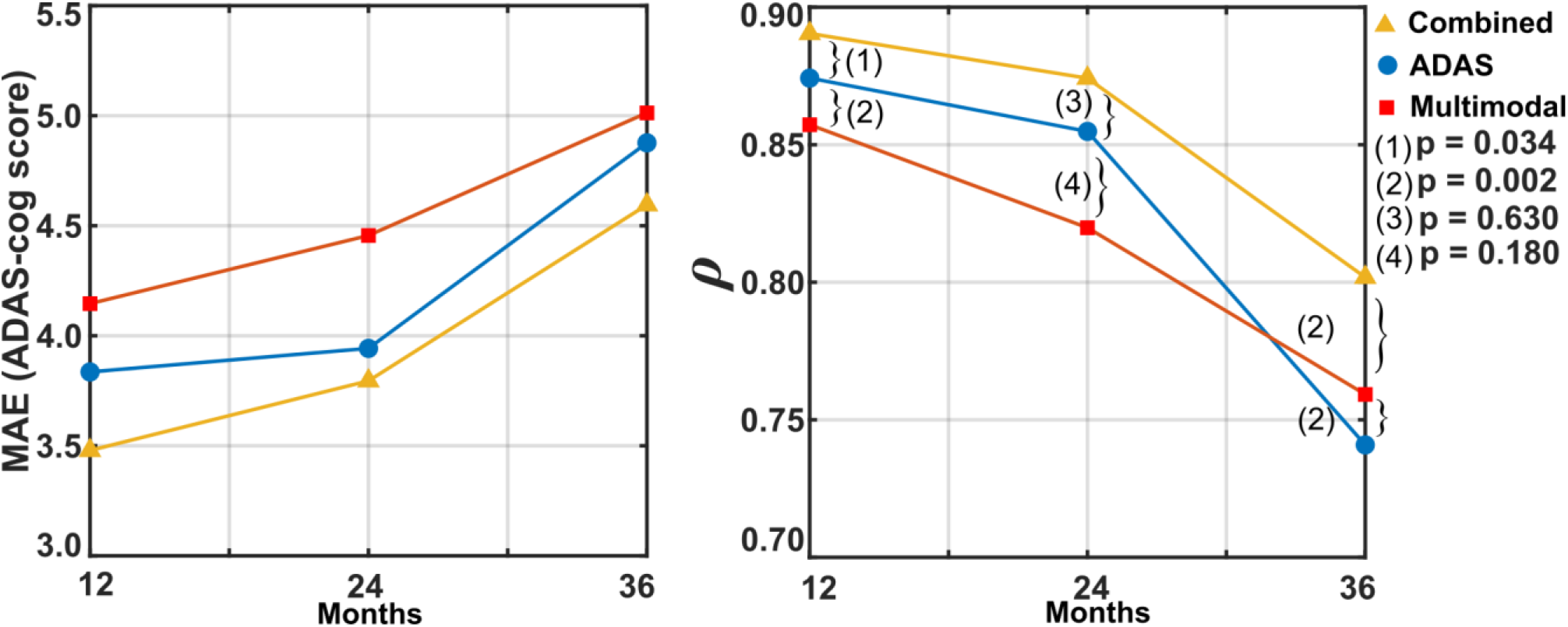
Performance comparison of prediction models utilizing only ADAS scores *vs*. multimodal data with and without the combination with baseline ADAS scores. The p-values correspond to pair-wise differences between the three prediction models at different time periods.

**Figure 6:**
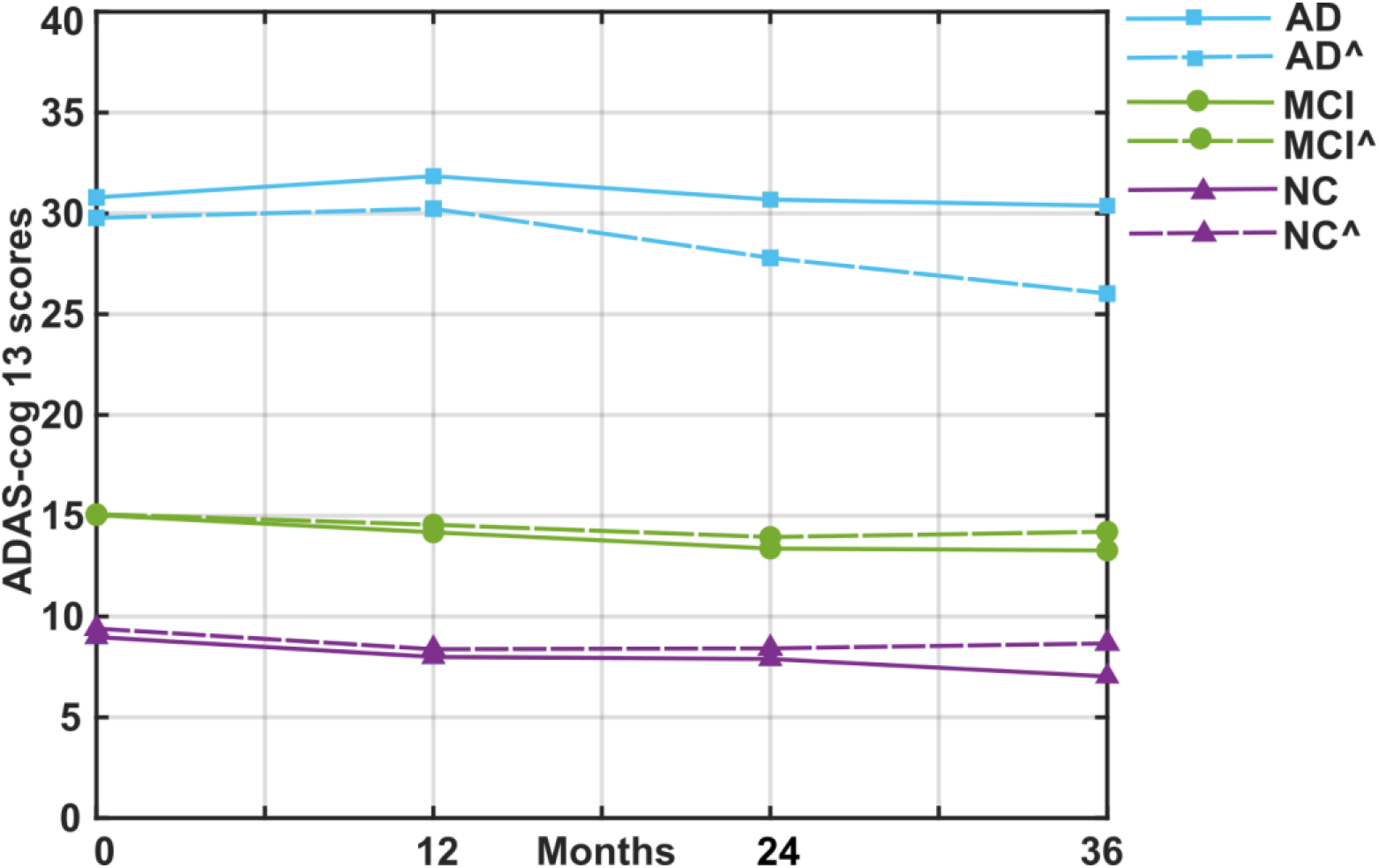
The mean of actual ADAS-cog 13 scores of subjects over 36 months is plotted (solid line) for the 3 diagnostic groups (normal cognition (NC), mild cognitive impairment (MCI) and Alzheimer’s disease (AD)). The learning model predicted scores for these time periods are also plotted (dashed lines) for each diagnostic group (model predictions indicated with a caret (^) symbol).

## 4 Discussion

We present a multi-modal regression approach to quantitatively track the progression of Alzheimer’s disease and show that it outperforms the conventional single modal approach. Quantification of AD aids clinicians in decisions with treatment and a multi-modal approach ensures that the prediction models consider all biomarkers contributing to the disease condition. Furthermore, conventional classification of patients into normal, MCI or AD could be avoided as a clear distinction amongst the group is a challenging task [52].

The classification of subjects based on a few modalities has been the focus of most recent studies. Although high classification accuracy (>80%) has been reported [11], we speculate that the impact of mislabeling a subject in the wrong category (and hence, wrong therapy prescribed) is higher than the error in predicting ADAS-cog scores (<5 units). Additionally, ADAS-cog scores are easy to interpret and follow the longitudinal tracking of AD progression. In agreement with the classification-based studies [4], the multi-modal approach outperforms the single modality, however, in this study multi-modal data were used for predicting the ADAS-cog scores. Furthermore, our multi-modal approach shows that ADAS-cog scores are conducive to longitudinal predictions contrary to Marinescu et al [21], where ADAS-cog scores were concluded not predictable. We, however, acknowledge that studies were not set up equally as there were time constraints, differences in subjects and underutilization of longitudinal data.

Clinically, NePB tests and ADAS-cog scores measure the subject’s cognitive abilities and this similarity was showcased with the observance of higher correlations (**Figure 3**). CSF biomarkers showed high correlations several studies support this strong relationship between CSF biomarkers and AD state [34]. As the precise pathophysiology and relative contribution of different pathogenic factors to AD at different phases of disease progression are currently still under investigation, the results advocate that instead of manually estimating the best markers, a multi-modal approach is beneficial. However, we acknowledge that the variable selection methods can be utilized to select the best AD measures (or create sparse models) utilized in multimodal modeling further improving the robustness of the prediction model.

The multivariate techniques (i.e., PLSR, SVR, and RF) were observed to perform very similarly in their predictions but the computation times were different, and this prompted us to favor PLSR. Other nonlinear model selection techniques could improve current results [53]. The subject attrition during follow-ups may have diminished the predictive performance of the model.

## 5 Conclusion

ADAS-cog 13 scores reflect the current cognitive state of individuals, and through multivariate regression and a multi-modal dataset, our results show that quantitative longitudinal prediction of AD progression is possible. Thus, the automated multi-modal approach may help clinicians make timely decisions for interventions at all stages of AD and inform likely disease progression at the start of clinical trials.

## Supporting information

Supplementary Section

## 7 Figure legends

### Figures in supplementary section

**Figure S.1**: The mean ADAS-cog 13 scores of subjects for different time periods and conversion of subjects in different categories. The subjects are grouped by the diagnosis as normal cognition (NC), mild cognitive impairment (MCI) and Alzheimer’s disease (AD).

**Figure S.2:** Regression modeling structure. Single modality uses one predictor at a time while multi-modal uses all the predictors as indicated above. The sample size for baseline (N = 757), 12months (N = 629), 24-months (N = 563) and 36-months (N = 314) were different due to missing values (cohort attrition). The predictors consist of age at baseline, years of formal education (Edu.), APOE e4 status (absence, single copy or homozygous coded as 0, 1 and 2 respectively), MRI-derived parameters, neuropsychiatric and behavioral assessment (NePB), AV45-PET measurements, CSF biomarkers (amyloid-β, τ, pτ) and FDG-PET measures. The number of features is indicated above each modality abbreviations. All the variables were considered as continuous and standardized to be zero-mean and unit standard deviation.

**Figure S.3:** Comparison of single and multi-modal dataset performance - collapsing across diagnostic status - with partial least squares regression (PLSR), support-vector regression (SVR) and random forest regression (RF). The performances are shown for cross-validation correlation (***ρ***) and mean absolute error (MAE).

**Figure S.4:** Genetic algorithm-based importance of parameters in contributing to increasing correlation as observed for 100 runs for the 36-month time period. The frequency indicates the proportional contribution in ADAS-cog 13 score prediction.

## 8 Table legends

**Table S.1:** Specific variable names from the TADPOLE D1_D2 dataset (1 to 5, details as in TADPOLE_D1_D2_Dict.csv) and FDG-PET from UCBERKELEYFDG_07_30_15 table.

**Table S.2:** Summary of the subject demographics and ADAS-cog 13 scores.

## Acknowledgments

This study was funded by the Research Committee of the Kuopio University Hospital Catchment Area for the State Research Funding (5041778) and The Finnish Foundation for Technology Promotion (8193, 6227) as well as The Academy of Finland (grant 316258 to JT).

Data collection and sharing for this project was funded by the Alzheimer’s Disease Neuroimaging Initiative (ADNI) (National Institutes of Health Grant U01 AG024904) and DOD ADNI (Department of Defense award number W81XWH-12-2-0012). ADNI is funded by the National Institute on Aging, the National Institute of Biomedical Imaging and Bioengineering, and through generous contributions from the following: AbbVie, Alzheimer’s Association; Alzheimer’s Drug Discovery Foundation; Araclon Biotech; BioClinica, Inc.; Biogen; Bristol-Myers Squibb Company; CereSpir, Inc.; Cogstate; Eisai Inc.; Elan Pharmaceuticals, Inc.; Eli Lilly and Company; EuroImmun; F. Hoffmann-La Roche Ltd and its affiliated company Genentech, Inc.; Fujirebio; GE Healthcare; IXICO Ltd.; Janssen Alzheimer Immunotherapy Research & Development, LLC.; Johnson & Johnson Pharmaceutical Research & Development LLC.; Lumosity; Lundbeck; Merck & Co., Inc.; Meso Scale Diagnostics, LLC.; NeuroRx Research; Neurotrack Technologies; Novartis Pharmaceuticals Corporation; Pfizer Inc.; Piramal Imaging; Servier; Takeda Pharmaceutical Company; and Transition Therapeutics. The Canadian Institutes of Health Research is providing funds to support ADNI clinical sites in Canada. Private sector contributions are facilitated by the Foundation for the National Institutes of Health (www.fnih.org). The grantee organization is the Northern California Institute for Research and Education, and the study is coordinated by the Alzheimer’s Therapeutic Research Institute at the University of Southern California. ADNI data are disseminated by the Laboratory for Neuro Imaging at the University of Southern California.

This work made use of the TADPOLE data sets https://tadpole.grand-challenge.org constructed by the EuroPOND consortium http://europond.eu funded by the European Union’s Horizon 2020 research and innovation program under grant agreement No 666992. The computational analysis was run on the servers provided by Bioinformatics Center, University of Eastern Finland, Finland.

## 9 Highlights

- A quantitative approach to track Alzheimer’s disease.
- The multi-modal approach enabled predicting ADAS-cog scores from 12 to 36 months.
- The combination of multimodal data and baseline ADAS scores enhanced the predictions of future timelines.

## 10 Authors contributions

**Prakash, M.:** Algorithm design and Data analysis.

**Abdelaziz, M.:** Data analysis.

**Zhang, L.** and **Strange, B.:** Clinical validation and supervision.

**Tohka, J.:** Study design and conception.

All authors contributed to the preparation and approval of the final submitted manuscript.

## References

[1] Gaudreault R, Mousseau N. Mitigating Alzheimer’s Disease with Natural Polyphenols: A Review. Curr Alzheimer Res 2019;16:529–43. https://doi.org/10.2174/1567205016666190315093520.

[2] Association A. 2019 Alzheimer’s disease facts and figures. Alzheimer’s Dement 2019;15:321–87. https://doi.org/10.1016/j.jalz.2019.01.010.

[3] Mueller SG, Weiner MW, Thal LJ, Petersen RC, Jack CR, Jagust W, et al. Ways toward an early diagnosis in Alzheimer’s disease: The Alzheimer’s Disease Neuroimaging Initiative (ADNI). Alzheimer’s Dement 2005;1:55–66. https://doi.org/10.1016/j.jalz.2005.06.003.

[4] Zhang D, Wang Y, Zhou L, Yuan H, Shen D. Multimodal classification of Alzheimer’s disease and mild cognitive impairment. Neuroimage 2011;55:856–67. https://doi.org/10.1016/j.neuroimage.2011.01.008.

[5] Perry RJ, Watson P, Hodges JR. The nature and staging of attention dysfunction in early (minimal and mild) Alzheimer’s disease: Relationship to episodic and semantic memory impairment. Neuropsychologia 2000;38:252–71. https://doi.org/10.1016/S0028-3932(99)00079-2.

[6] Morris JC, Storandt M, Miller JP, McKeel DW, Price JL, Rubin EH, et al. Mild cognitive impairment represents early-stage Alzheimer disease. Arch Neurol 2001;58:397–405. https://doi.org/10.1001/archneur.58.3.397.

[7] Nestor PJ, Scheltens P, Hodges JR. Advances in the early detection of alzheimer’s disease. Nat Rev Neurosci 2004;10:S34. https://doi.org/10.1038/nrn1433.

[8] Duara R, Barker WW, Lopez-Alberola R, Loewenstein DA, Grau LB, Gilchrist D, et al. Alzheimer’s disease: Interaction of apolipoprotein E genotype, family history of dementia, gender, education, ethnicity, and age of onset. Neurology 1996;46:1575–9. https://doi.org/10.1212/WNL.46.6.1575.

[9] Selkoe DJ. The molecular pathology of Alzheimer’s disease. Neuron 1991;6:487–98. https://doi.org/10.1016/0896-6273(91)90052-2.

[10] Teipel S, Drzezga A, Grothe MJ, Barthel H, Chételat G, Schuff N, et al. Multimodal imaging in Alzheimer’s disease: Validity and usefulness for early detection. Lancet Neurol 2015;14:1037–53. https://doi.org/10.1016/S1474-4422(15)00093-9.

[11] Rathore S, Habes M, Iftikhar MA, Shacklett A, Davatzikos C. A review on neuroimaging-based classification studies and associated feature extraction methods for Alzheimer’s disease and its prodromal stages. Neuroimage 2017;155:530–48. https://doi.org/10.1016/j.neuroimage.2017.03.057.

[12] Jack CR, Knopman DS, Jagust WJ, Petersen RC, Weiner MW, Aisen PS, et al. Tracking pathophysiological processes in Alzheimer’s disease: An updated hypothetical model of dynamic biomarkers. Lancet Neurol 2013;12:207–16. https://doi.org/10.1016/S1474-4422(12)70291-0.

[13] Keihaninejad S, Zhang H, Ryan NS, Malone IB, Modat M, Cardoso MJ, et al. An unbiased longitudinal analysis framework for tracking white matter changes using diffusion tensor imaging with application to Alzheimer’s disease. Neuroimage 2013;72:153–63. https://doi.org/10.1016/j.neuroimage.2013.01.044.

[14] Jack CR, Holtzman DM. Biomarker modeling of alzheimer’s disease. Neuron 2013;80:1347–58. https://doi.org/10.1016/j.neuron.2013.12.003.

[15] Growdon JH. Incorporating biomarkers into clinical drug trials in Alzheimer’s disease. J Alzheimers Dis 2001;3:287–92. https://doi.org/10.3233/jad-2001-3303.

[16] Yang E, Farnum M, Lobanov V, Schultz T, Raghavan N, Samtani MN, et al. Quantifying the pathophysiological timeline of Alzheimer’s disease. J Alzheimer’s Dis 2011;26:745–53. https://doi.org/10.3233/JAD-2011-110551.

[17] William-Faltaos D, Chen Y, Wang Y, Gobburu J, Zhu H. Quantification of disease progression and dropout for Alzheimer’s disease. Int J Clin Pharmacol Ther 2013;51:120–31. https://doi.org/10.5414/CP201787.

[18] Skinner J, Carvalho JO, Potter GG, Thames A, Zelinski E, Crane PK, et al. The Alzheimer’s Disease Assessment Scale-Cognitive-Plus (ADAS-Cog-Plus): An expansion of the ADAS-Cog to improve responsiveness in MCI. Brain Imaging Behav 2012;6:489–501. https://doi.org/10.1007/s11682-012-9166-3.

[19] Kueper JK, Speechley M, Montero-Odasso M. The Alzheimer’s Disease Assessment Scale-Cognitive Subscale (ADAS-Cog): Modifications and Responsiveness in Pre-Dementia Populations. A Narrative Review. J Alzheimer’s Dis 2018;63:423–44. https://doi.org/10.3233/JAD-170991.

[20] Zhang D, Shen D. Predicting Future Clinical Changes of MCI Patients Using Longitudinal and Multimodal Biomarkers. PLoS One 2012;7:e33182. https://doi.org/10.1371/journal.pone.0033182.

[21] Marinescu R V., Oxtoby NP, Young AL, Bron EE, Toga AW, Weiner MW, et al. TADPOLE Challenge: Accurate Alzheimer’s Disease Prediction Through Crowdsourced Forecasting of Future Data, 2019, p. 1–10. https://doi.org/10.1007/978-3-030-32281-6_1.

[22] Steenland K, Zhao L, Goldstein F, Cellar J, Lah J. Biomarkers for predicting cognitive decline in those with normal cognition. J Alzheimers Dis 2014;40:587–94. https://doi.org/10.3233/JAD-2014-131343.

[23] Benge JF, Balsis S, Geraci L, Massman PJ, Doody RS. How well do the ADAS-cog and its subscales measure cognitive dysfunction in Alzheimer’s disease? Dement Geriatr Cogn Disord 2009;28:63–9. https://doi.org/10.1159/000230709.

[24] Marinescu R V., Oxtoby NP, Young AL, Bron EE, Toga AW, Weiner MW, et al. TADPOLE Challenge: Prediction of Longitudinal Evolution in Alzheimer’s Disease 2018.

[25] Gómez-Sancho M, Tohka J, Gómez-Verdejo V. Comparison of feature representations in MRI-based MCI-to-AD conversion prediction. Magn Reson Imaging 2018;50:84–95. https://doi.org/10.1016/j.mri.2018.03.003.

[26] Voevodskaya O. The effects of intracranial volume adjustment approaches on multiple regional MRI volumes in healthy aging and Alzheimer’s disease. Front Aging Neurosci 2014;6. https://doi.org/10.3389/fnagi.2014.00264.

[27] Reuter M, Schmansky NJ, Rosas HD, Fischl B. Within-subject template estimation for unbiased longitudinal image analysis. Neuroimage 2012;61:1402–18. https://doi.org/10.1016/j.neuroimage.2012.02.084.

[28] Johnson KA, Sperling RA, Gidicsin CM, Carmasin JS, Maye JE, Coleman RE, et al. Florbetapir (F18-AV-45) PET to assess amyloid burden in Alzheimer’s disease dementia, mild cognitive impairment, and normal aging. Alzheimer’s Dement 2013;9. https://doi.org/10.1016/j.jalz.2012.10.007.

[29] Landau SM, Mintun MA, Joshi AD, Koeppe RA, Petersen RC, Aisen PS, et al. Amyloid deposition, hypometabolism, and longitudinal cognitive decline. Ann Neurol 2012;72:578–86. https://doi.org/10.1002/ana.23650.

[30] Landau SM, Lu M, Joshi AD, Pontecorvo M, Mintun MA, Trojanowski JQ, et al. Comparing positron emission tomography imaging and cerebrospinal fluid measurements of β-amyloid. Ann Neurol 2013;74:826–36. https://doi.org/10.1002/ana.23908.

[31] Mormino EC, Kluth JT, Madison CM, Rabinovici GD, Baker SL, Miller BL, et al. Episodic memory loss is related to hippocampal-mediated β-amyloid deposition in elderly subjects. Brain 2009;132:1310–23. https://doi.org/10.1093/brain/awn320.

[32] Landau SM, Harvey D, Madison CM, Koeppe RA, Reiman EM, Foster NL, et al. Associations between cognitive, functional, and FDG-PET measures of decline in AD and MCI. Neurobiol Aging 2011;32:1207–18. https://doi.org/10.1016/j.neurobiolaging.2009.07.002.

[33] Jagust WJ, Bandy D, Chen K, Foster NL, Landau SM, Mathis CA, et al. The Alzheimer’s Disease Neuroimaging Initiative positron emission tomography core. Alzheimer’s Dement 2010;6:221–9. https://doi.org/10.1016/j.jalz.2010.03.003.

[34] Shaw LM, Vanderstichele H, Knapik-Czajka M, Clark CM, Aisen PS, Petersen RC, et al. Cerebrospinal fluid biomarker signature in alzheimer’s disease neuroimaging initiative subjects. Ann Neurol 2009;65:403–13. https://doi.org/10.1002/ana.21610.

[35] Battista P, Salvatore C, Castiglioni I. Optimizing neuropsychological assessments for cognitive, behavioral, and functional impairment classification: A machine learning study. Behav Neurol 2017;2017. https://doi.org/10.1155/2017/1850909.

[36] Folstein MF, Folstein SE, McHugh PR. “Mini-mental state”. A practical method for grading the cognitive state of patients for the clinician. J Psychiatr Res 1975;12:189–98. https://doi.org/10.1016/0022-3956(75)90026-6.

[37] Rey A. L’examin clinique en psychologie. Press Univ Fr 1958. https://psycnet.apa.org/record/1959-03776-000 (accessed December 30, 2019).

[38] Pfeffer RI, Kurosaki TT, Harrah CH, Chance JM, Filos S. Measurement of functional activities in older adults in the community. Journals Gerontol 1982;37:323–9. https://doi.org/10.1093/geronj/37.3.323.

[39] Prencipe M, Casini AR, Ferretti C, Lattanzio MT, Fiorelli M, Culasso F. Prevalence of dementia in an elderly rural population: effects of age, sex, and education. J Neurol Neurosurg Psychiatry 1996;60:628–33. https://doi.org/10.1136/jnnp.60.6.628.

[40] Michaelson DM. APOE ε4: The most prevalent yet understudied risk factor for Alzheimer’s disease. Alzheimer’s Dement 2014;10:861–8. https://doi.org/10.1016/j.jalz.2014.06.015.

[41] Mohs RC, Knopman D, Petersen RC, Ferris SH, Ernesto C, Grundman M, et al. Development of cognitive instruments for use in clinical trials of antidementia drugs: Additions to the Alzheimer’s disease assessment scale that broaden its scope. Alzheimer Dis Assoc Disord 1997;11. https://doi.org/10.1097/00002093-199700112-00003.

[42] Raghavan N, Samtani MN, Farnum M, Yang E, Novak G, Grundman M, et al. The ADAS-Cog revisited: Novel composite scales based on ADAS-Cog to improve efficiency in MCI and early AD trials. Alzheimer’s Dement 2013;9. https://doi.org/10.1016/j.jalz.2012.05.2187.

[43] Krishnan A, Williams LJ, McIntosh AR, Abdi H. Partial Least Squares (PLS) methods for neuroimaging: A tutorial and review. Neuroimage 2011;56:455–75. https://doi.org/10.1016/j.neuroimage.2010.07.034.

[44] Zhang D, Shen D. Multi-modal multi-task learning for joint prediction of multiple regression and classification variables in Alzheimer’s disease. Neuroimage 2012;59:895–907. https://doi.org/10.1016/j.neuroimage.2011.09.069.

[45] Gray KR, Aljabar P, Heckemann RA, Hammers A, Rueckert D. Random forest-based similarity measures for multi-modal classification of Alzheimer’s disease. Neuroimage 2013;65:167–75. https://doi.org/10.1016/j.neuroimage.2012.09.065.

[46] Leardi R, Boggia R, Terrile M. Genetic algorithms as a strategy for feature selection. J Chemom 1992;6:267–81. https://doi.org/10.1002/cem.1180060506.

[47] Lewis JD, Evans AC, Tohka J. T1 white/gray contrast as a predictor of chronological age, and an index of cognitive performance. Neuroimage 2018;173:341–50. https://doi.org/10.1016/j.neuroimage.2018.02.050.

[48] de Jong S. SIMPLS: An alternative approach to partial least squares regression. Chemom Intell Lab Syst 1993;18:251–63. https://doi.org/10.1016/0169-7439(93)85002-X.

[49] Breiman L. Bagging predictors. Mach Learn 1996;24:123–40. https://doi.org/10.1007/BF00058655.

[50] Breiman L. Random forests. Mach Learn 2001;45:5–32. https://doi.org/10.1023/A:1010933404324.

[51] R. Leardi and A. Lupiáñez. Genetic algorithms applied to feature selection in PLS regression: how and when to use them. Chemom Intell Lab Syst 1998;41:95–207.

[52] Leyhe T, Reynolds CF, Melcher T, Linnemann C, Klöppel S, Blennow K, et al. A common challenge in older adults: Classification, overlap, and therapy of depression and dementia. Alzheimer’s Dement 2017;13:59–71. https://doi.org/10.1016/j.jalz.2016.08.007.

[53] Chouldechova A, Hastie T. Generalized Additive Model Selection. ArXiv Prepr ArXiv 150603850 2015.

