## Supplementary Section for "Quantitative longitudinal predictions of Alzheimer’s disease by multi-modal predictive learning"

### Supplementary Material

#### A. Methods

The variable names of the modalities are provided in Table A.1.

**Table S.1:** Specific variable names from the TADPOLE D1_D2 dataset (1 to 5, details as in TADPOLE_D1_D2_Dict.csv) and FDG-PET from UCBERKELEYFDG_07_30_15 table.

| | Category | Variable name |
|---|---|---|
| 1 | Subject details | PTEDUCAT |
| | | APOE4 |
| | | AGE |
| 2 | AVF45-PET | FRONTAL_UCBERKELEYAV45_10_17_16 |
| | | CINGULATE_UCBERKELEYAV45_10_17_16 |
| | | PARIETAL_UCBERKELEYAV45_10_17_16 |
| | | TEMPORAL_UCBERKELEYAV45_10_17_16 |
| 3 | MRI parameters | Ventricles |
| | | Hippocampus |
| | | WholeBrain |
| | | Entorhinal |
| | | Fusiform |
| | | MidTemp |
| | | ICV |
| 4 | Cognitive measures | MMSE |
| | | RAVLT_learning |
| | | RAVLT_immediate |
| | | RAVLT_perc_forgetting |
| | | FAQ |
| 5 | CSF | ABETA_UPENNBIOMK9_04_19_17 |
| | | TAU_UPENNBIOMK9_04_19_17 |
| | | PTAU_UPENNBIOMK9_04_19_17 |
| 6 | FDG-PET | Angular left |
| | | Angular right |
| | | CingulumPost |
| | | Temporal left |
| | | Temporal right |

**Regression methods:**

A brief background of the methods used in this study is presented.

**Partial least squares regression (PLSR)**

The difference between PLSR and ordinary least squares (OLS) lies in the fact that PLSR is essentially a dimensionality reduction method in which the initial set of features $X_i$, (i = 0, 1, .., L) are mapped to a new set of features $Z_k$, (k = 0, 1, .., M), where M < L and k are optimized with the training data. This property is a very useful one especially when the number of available data samples is not necessarily large enough compared to the number of features, which is the case in our problem. The new set of features are obtained in a supervised manner in which the response Y is used together with the old features $X_i$ to identify the new features $Z_k$ [1].

**Support vector regression (SVR)**

Instead of minimizing the sum of squared errors between the model output and the actual response as in OLS regression, SVR aims at minimizing a different type of cost function, where only residuals larger in absolute value than some margin of tolerance contribute to the cost function. Towards this end, a regularization parameter is also defined to indicate the amount of penalty applied to data samples lying beyond the margin of tolerance. This regularization parameter represents the first SVR hyperparameter. Additionally, SVR provides the option to map the input data points using linear or nonlinear kernels (or basis functions) that can provide better prediction performance, especially when there is some nonlinear dependency between the ADAS scores and the set of features.

**Random forest regression (RF)**

Random forest methods [2], unlike PLSR or SVR, utilize a combination of results of many models/trees (ensemble of models). Trees are constructed via classification and regression tree (CART) and the data is sampled via bagging and the average of the outputs used for classification or regression modeling. A comprehensive review of RF on neuroimaging data is provided by Sarica et al [3].

**A genetic algorithm for variable importance**

The selection of the best subset of variables, termed variable or feature selection, is a challenging problem in machine learning. One of the various approaches for variable selection is by Genetic algorithms (GA) [4], GA-based variable selection can provide us a ranking of importance of individual variables for ADAS prediction. GA aims to select a subset of variables that best predict the response variable, ADAS-cog score in our case. Due to randomness inherent to the GA, a different set of variables is selected during each run of the GA. By running GA r (here r = 100) times, we assume that the frequency of retention of a variable among r GA runs corresponds to the importance of that variable in predicting the ADAS score.

GA is a general optimization technique inspired by natural selection. GA-based variable selection codes each possible solution to the problem as a binary vector with the same number of components as there are variables, with 1 (0) in the $i^{th}$ component meaning that the ith variable is (not) in the model.  GA initializes with a random population of potential solutions (called chromosomes) and then evolves this population through mutation, selection and cross-over operators guided by the fitness function, which, in this case, is a cross-validated mean squared error between the true ADAS-cog score and predicted ADAS-cog score.

We employed the GA-PLS toolbox by Leardi et al to create PLS models [5]. The *gapls* function from the toolbox was used and the frequency of retention of variables in the models was recorded with default settings [5]. The GA internally employs cross-validation and root mean square error (RMSECV) on PLS models to detect the change in this error and decide to either retain, delete, and/or carry over a subset of variables to the next iteration.

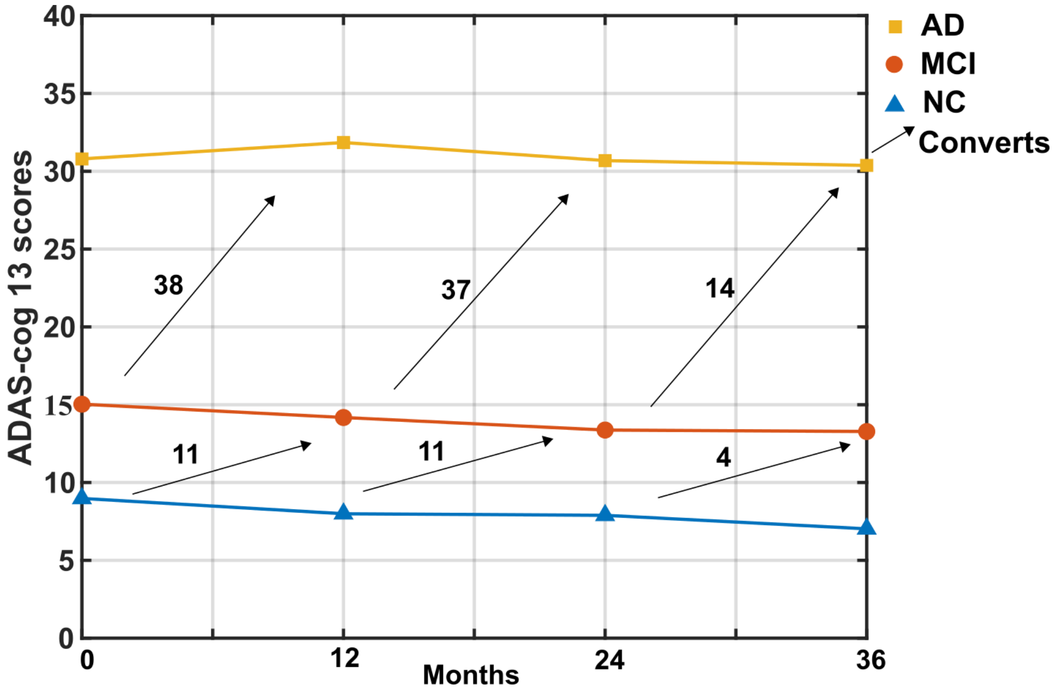


**Figure S.1**: The mean ADAS-cog 13 scores of subjects for different time periods and conversion of subjects in different categories. The subjects are grouped by the diagnosis as normal cognition (NC), mild cognitive impairment (MCI) and Alzheimer's disease (AD).

**Table S.2:** Summary of the subject demographics and ADAS-cog 13 scores.

| | Baseline | | | 12 months later | | | 24 months later | | | 36 months later | | |
|---|---|---|---|---|---|---|---|---|---|---|---|---|
| | **NC** | **MCI** | **AD** | **NC** | **MCI** | **AD** | **NC** | **MCI** | **AD** | **NC** | **MCI** | **AD** |
| **No. of Subjects** | 241 | 405 | 111 | 169 | 337 | 117 | 195 | 271 | 89 | 40 | 209 | 61 |
| **M/F** | 112/129 | 226/179 | 68/43 | 89/80 | 181/156 | 76/41 | 103/92 | 147/124 | 53/36 | 22/18 | 101/108 | 39/22 |
| **Age** | 56-89 | 55-91 | 55-90 | 55-89 | 56-92 | 55-90 | 56-89 | 55-91 | 55-90 | 55-84 | 55-87 | 55-56 |
| **ADAS-cog 13 scores** | 0-24 | 3-38 | 14-51 | 1-21 | 1-73 | 9-62 | 0-25 | 2-45 | 8-67 | 0-18 | 0-36 | 8-74 |

**RIDS of subjects**:

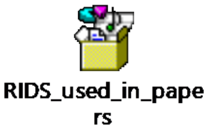


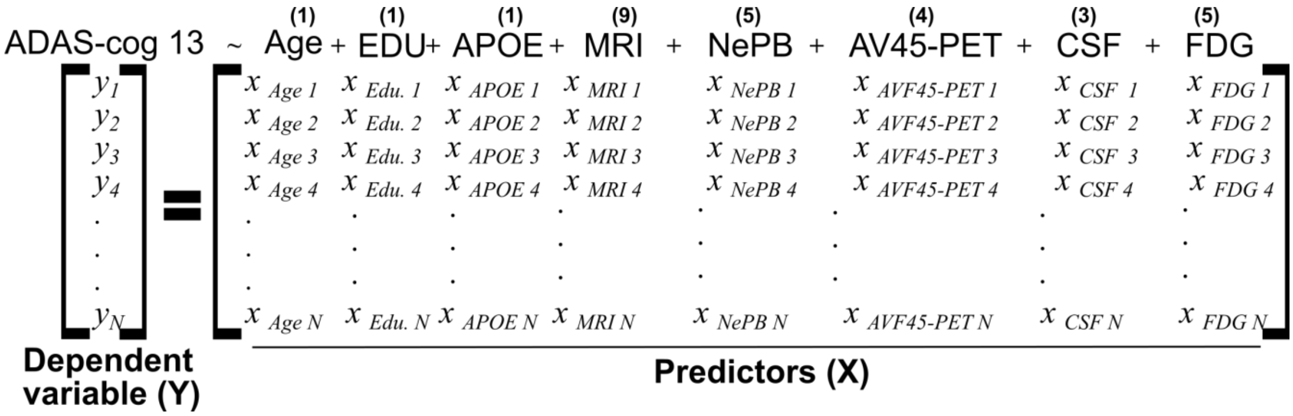


**Figure S.2:** Regression modeling structure**.** Single modality uses one predictor at a time while multi-modal uses all the predictors as indicated above. The sample size for baseline (N = 757), 12-months (N = 629), 24-months (N = 563) and 36-months (N = 314) were different due to missing values (cohort attrition). The predictors consist of age at baseline, years of formal education (Edu.), APOE e4 status (absence, single copy or homozygous coded as 0, 1 and 2 respectively), MRI-derived parameters, neuropsychiatric and behavioral assessment (NePB), AV45-PET measurements, CSF biomarkers (amyloid-β, τ, pτ) and FDG-PET measures. The number of features is indicated above each modality abbreviations. All the variables were considered as continuous and standardized to be zero-mean and unit standard deviation.

### B. Brief description of the structural MRI data processing

In this section, we provide a brief description of the processing performed in the manuscript on the baseline MRI data which was downloaded from the ADNI web site (www.loni.ucla.edu/ADNI). Cortical reconstruction and volumetric segmentation are performed with the FreeSurfer 5.1 image analysis suite, which is documented and freely available for download online (http://surfer.nmr.mgh.harvard.edu/).

Briefly, this processing includes motion correction and averaging [6] of multiple volumetric T1 weighted images (when more than one is available), removal of non-brain tissue using a hybrid watershed/surface deformation procedure [7], automated Talairach transformation, segmentation of the subcortical white matter and deep gray matter volumetric structures (including hippocampus, amygdala, caudate, putamen, ventricles) [8,9] intensity normalization [10], tessellation of the gray matter white matter boundary, automated topology correction [11,12], and surface deformation following intensity gradients to optimally place the gray/white and gray/cerebrospinal fluid borders at the location where the greatest shift in intensity defines the transition to the other tissue class [13,14]. Once the cortical models are complete, a number of deformable procedures can be performed for further data processing and analysis including surface inflation, registration to a spherical atlas which is based on individual cortical folding patterns to match cortical geometry across subjects [15], parcellation of the cerebral cortex into units with respect to gyral and sulcal structure [8], and creation of a variety of surface-based data including maps of curvature and sulcal depth. This method uses both intensity and continuity information from the entire three dimensional MR volume in segmentation and deformation procedures to produce representations of cortical thickness, calculated as the closest distance from the gray/white boundary to the gray/CSF boundary at each vertex on the tessellated surface [14]. The maps are created using spatial intensity gradients across tissue classes and are therefore not simply reliant on absolute signal intensity. The maps produced are not restricted to the voxel resolution of the original data thus are capable of detecting submillimeter differences between groups. Procedures for the measurement of cortical thickness have been validated against histological analysis [16] and manual measurements [17,18]. Freesurfer morphometric procedures have been demonstrated to show good test-retest reliability across scanner manufacturers and across field strengths [19].

### C. Results

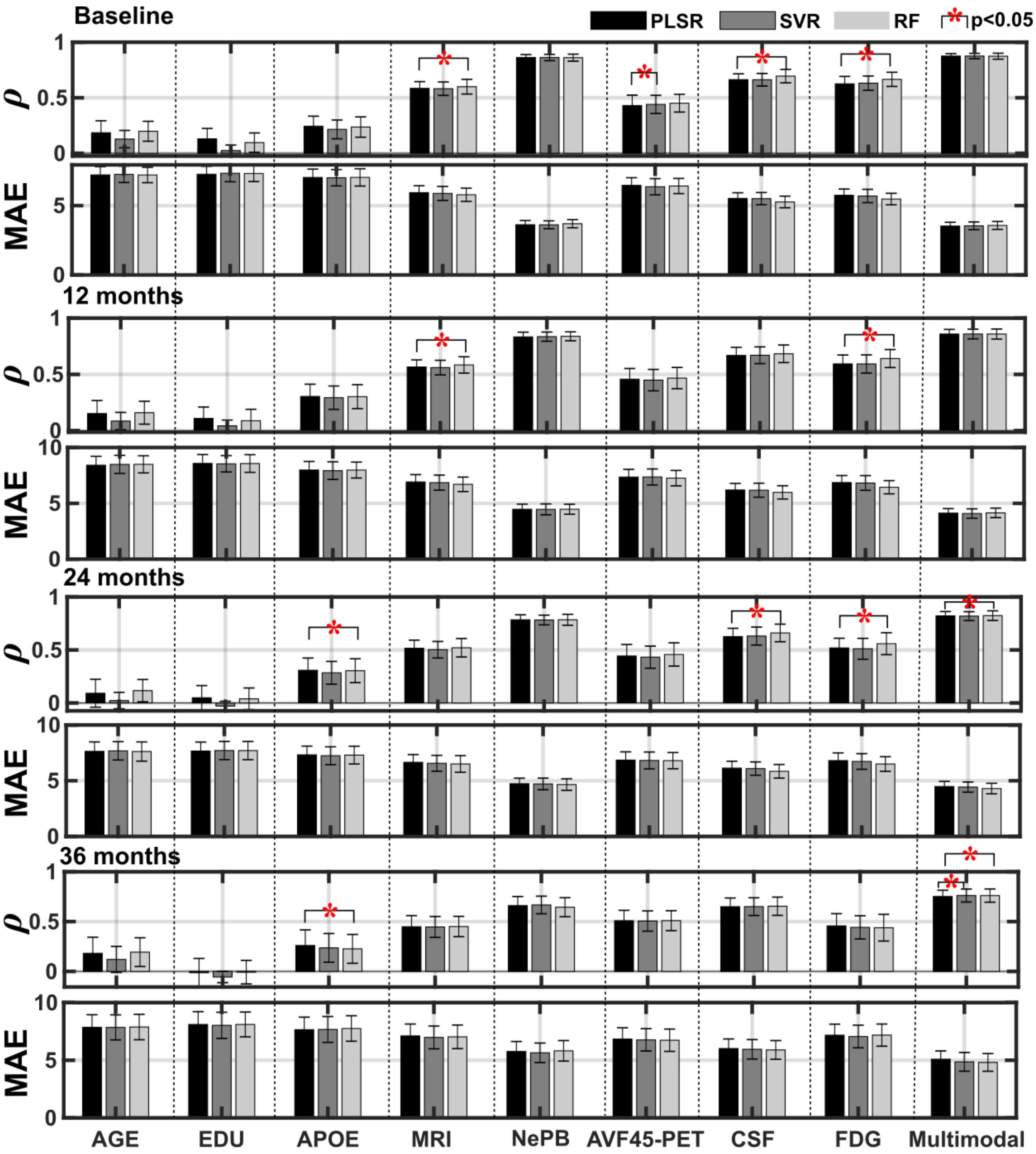


**Figure S.3:** Comparison of single and multi-modal dataset performance – collapsing across diagnostic status – with partial least squares regression (PLSR), support-vector regression (SVR) and random forest regression (RF). The performances are shown for cross-validation correlation (***ρ***) and mean absolute error (MAE).

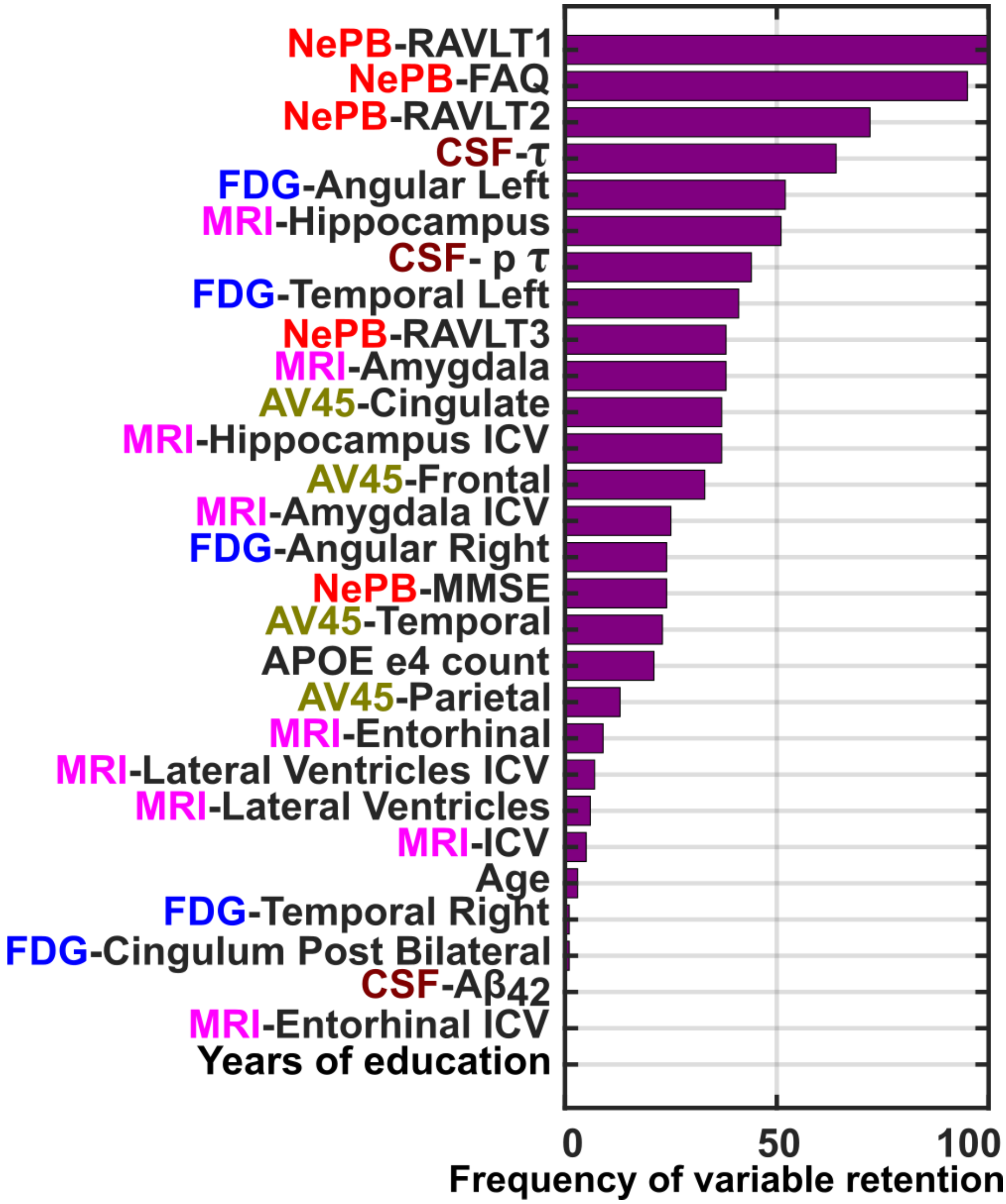


**Figure S.4:** Genetic algorithm-based importance of parameters in contributing to increasing correlation as observed for 100 runs for the 36-month time period. The frequency indicates the proportional contribution in ADAS-cog 13 score prediction.

**References**:


[1] Krishnan A, Williams LJ, McIntosh AR, Abdi H. Partial Least Squares (PLS) methods for neuroimaging: A tutorial and review. Neuroimage 2011;56:455–75. https://doi.org/10.1016/j.neuroimage.2010.07.034.

[2] Breiman L. Random forests. Mach Learn 2001;45:5–32. https://doi.org/10.1023/A:1010933404324.

[3] Sarica A, Cerasa A, Quattrone A. Random forest algorithm for the classification of neuroimaging data in Alzheimer’s disease: A systematic review. Front Aging Neurosci 2017;9. https://doi.org/10.3389/fnagi.2017.00329.

[4] Leardi R, Boggia R, Terrile M. Genetic algorithms as a strategy for feature selection. J Chemom 1992;6:267–81. https://doi.org/10.1002/cem.1180060506.

[5] R. Leardi and A. Lupiáñez. Genetic algorithms applied to feature selection in PLS regression: how and when to use them. Chemom Intell Lab Syst 1998;41:95–207.

[6] Reuter M, Rosas HD, Fischl B. Highly accurate inverse consistent registration: A robust approach. Neuroimage 2010;53:1181–96. https://doi.org/10.1016/j.neuroimage.2010.07.020.

[7] Ségonne F, Dale AM, Busa E, Glessner M, Salat D, Hahn HK, et al. A hybrid approach to the skull stripping problem in MRI. Neuroimage 2004;22:1060–75. https://doi.org/10.1016/j.neuroimage.2004.03.032.

[8] Fischl B, Salat DH, Busa E, Albert M, Dieterich M, Haselgrove C, et al. Whole brain segmentation: Automated labeling of neuroanatomical structures in the human brain. Neuron 2002;33:341–55. https://doi.org/10.1016/S0896-6273(02)00569-X.

[9] Fischl B, Salat DH, Van Der Kouwe AJW, Makris N, Ségonne F, Quinn BT, et al. Sequence-independent segmentation of magnetic resonance images. Neuroimage, vol. 23, 2004. https://doi.org/10.1016/j.neuroimage.2004.07.016.

[10] Sied JG, Zijdenbos AP, Evans AC. A nonparametric method for automatic correction of intensity nonuniformity in mri data. IEEE Trans Med Imaging 1998;17:87–97.

https://doi.org/10.1109/42.668698.

[11] Fischl B, Liu A, Dale AM. Automated manifold surgery: Constructing geometrically accurate and topologically correct models of the human cerebral cortex. IEEE Trans Med Imaging 2001;20:70–80. https://doi.org/10.1109/42.906426.

[12] Ségonne F, Pacheco J, Fischl B. Geometrically accurate topology-correction of cortical surfaces using nonseparating loops. IEEE Trans Med Imaging 2007;26:518–29. https://doi.org/10.1109/TMI.2006.887364.

[13] Dale AM, Fischl B, Sereno MI. Cortical surface-based analysis: I. Segmentation and surface reconstruction. Neuroimage 1999;9:179–94. https://doi.org/10.1006/nimg.1998.0395.

[14] Fischl B, Dale AM. Measuring the thickness of the human cerebral cortex from magnetic resonance images. Proc Natl Acad Sci U S A 2000;97:11050–5. https://doi.org/10.1073/pnas.200033797.

[15] Fischl B, Sereno MI, Tootell RBH, Dale AM. High-resolution intersubject averaging and a coordinate system for the cortical surface. Hum Brain Mapp 1999;8:272–84. https://doi.org/10.1002/(SICI)1097-0193(1999)8:4<272::AID-HBM10>3.0.CO;2-4.

[16] Rosas HD, Liu AK, Hersch S, Glessner M, Ferrante RJ, Salat DH, et al. Regional and progressive thinning of the cortical ribbon in Huntington's disease. Neurology 2002;58:695–701. https://doi.org/10.1212/WNL.58.5.695.

[17] Kuperberg GR, Broome MR, McGuire PK, David AS, Eddy M, Ozawa F, et al. Regionally localized thinning of the cerebral cortex in schizophrenia. Arch Gen Psychiatry 2003;60:878–88. https://doi.org/10.1001/archpsyc.60.9.878.

[18] Salat DH, Buckner RL, Snyder AZ, Greve DN, Desikan RSR, Busa E, et al. Thinning of the cerebral cortex in aging. Cereb Cortex 2004;14:721–30. https://doi.org/10.1093/cercor/bhh032.

[19] Reuter M, Schmansky NJ, Rosas HD, Fischl B. Within-subject template estimation for

unbiased longitudinal image analysis. Neuroimage 2012;61:1402–18. https://doi.org/10.1016/j.neuroimage.2012.02.084.
